# Adaptive Neural Reorganization Enables Real-Time Finger-Level Robotic Control in BCI-Naïve Stroke Survivors

**DOI:** 10.64898/2026.06.15.732267

**Authors:** Yidan Ding, Maxim Karrenbach, Zachary Johnson, Hanwen Wang, Jintao Zhang, George F. Wittenberg, Bin He

## Abstract

Restoring hand function remains a major challenge for individuals with motor impairments following stroke. Noninvasive brain–computer interfaces (BCIs) aim to address this problem by translating neural signals into robotic assistance; however, control of individual fingers has not been demonstrated in BCI-naïve populations. In this study, we investigated whether individuals with stroke and no prior BCI experience could achieve finger-level robotic control using motor imagery. Nine stroke-affected participants performed real-time BCI tasks to control a robotic hand through imagined finger movements decoded from electroencephalography. On average, participants achieved decoding accuracies of 84% for two-finger tasks and 61% for three-finger tasks, demonstrating reliable control at the level of individual fingers. These results indicate that discriminable neural signals for fine motor control persist after stroke and can be leveraged using data-driven deep learning decoders. Sensor-level and source-level electrophysiological analyses further reveal patterns of stroke-related neural reorganization. Overall, these findings support the potential of noninvasive, finger-level BCIs for post-stroke robotic assistance.

## Introduction

Stroke remains one of the leading causes of long-term motor disability worldwide, with upper-limb impairment among the most common and debilitating consequences, affecting nearly half of all stroke survivors [1]. Restoration of hand and arm function is consistently ranked as a top priority by both patients and researchers, as it directly impacts independence in daily activities and overall quality of life [2–4]. In response, neural engineering efforts have increasingly focused on restoring biological limb function and enabling robotic control through neuroprosthetic approaches and neuromodulatory interventions [5–11].

Noninvasive brain–computer interfaces (BCIs) have emerged as a promising approach to augment motor recovery and functional compensation by translating neural activity into control signals, thereby bypassing damaged neuromuscular pathways [12–15]. Their noninvasive nature makes them particularly suitable for widespread adoption among stroke population, as well as the broader population living with paralysis. By linking cortical signals associated with movement intention to real-time feedback, BCIs may facilitate adaptive neuroplasticity while also providing functional assistance. Over the past two decades, upper-limb BCI systems have demonstrated their potential to restore gross motor functions through robotic assistance [16–21] and to promote neural recovery using functional electrical stimulation as an effector [22,23].

Nevertheless, BCI applications targeting stroke survivors have largely been confined to coarse, limb-level control —such as whole-hand opening/closing or wrist flexion—rather than fine, dexterous movements. This limitation arises from both technological and neurophysiological challenges. From a signal processing perspective, decoding naturalistic, multi-finger movements from noninvasive EEG is difficult due to low spatial resolution caused by volume conduction [24] and overlapping cortical representations of individual digits [25]. From a clinical perspective, stroke-induced lesions and brain network disruptions introduce substantial variability in neural activity, complicating the extraction of reliable motor intention signals [20,26,27]. Consequently, it remains unclear whether individuals with stroke-related impairments can achieve reliable, independent finger-level control using noninvasive BCIs.

Recent work in BCI-experienced healthy participants has demonstrated that precise, naturalistic robotic hand control at the individual finger level is achievable using electroencephalography (EEG)-based decoding, combined with real-time feedback and adaptive training [28]. Building upon these findings, we hypothesize that BCI-naïve stroke survivors—despite altered cortical oscillatory dynamics and connectivity—can also learn to achieve reliable finger-specific BCI control through effective online training and intuitive task design. Specifically, we propose that data-driven decoding models can identify compensatory spectral–spatial features that bypass lesioned regions and leverage preserved motor representations.

To test this hypothesis, we developed a multi-session, online BCI training protocol in which stroke survivors performed individual-finger motor imagery tasks with concurrent robotic feedback. This study aims to evaluate the feasibility and learning dynamics of fine-grained finger control using EEG-based BCIs in stroke populations, and to characterize how stroke-related neural reorganization influences decoding performance and feature representations. Collectively, these findings provide insight into the adaptability of post-stroke motor networks and lay the groundwork for next-generation neuroprosthetic and rehabilitation BCIs capable of restoring dexterous hand function.

## Results

In this study, we investigated the feasibility of EEG-based, finger-level, real-time robotic hand control in BCI-naïve stroke-affected individuals (n = 9; Table 1), including six participants with unilateral motor impairment. Participants first completed two motor imagery (MI) training sessions involving a one-dimensional (1D) cursor control task to familiarize them with limb-level MI. They then participated in two offline sessions followed by seven online sessions of unilateral finger MI tasks, using the same experimental paradigm as described in Ding et al. [28]. Participants without motor impairment (n = 3) performed finger MI using their dominant hand, whereas those with motor deficits (n = 6) performed finger MI using their affected hand. The offline sessions were used for task familiarization and subject-specific decoder training. During the online BCI sessions, an EEGNet-8,2 model [29] was used for real-time decoding of imagined finger movements to control a robotic hand. EEG signals were band-pass filtered between 4–40 Hz for the first five online sessions and between 0.5–40 Hz for the final two sessions. Each online session consisted of 32 runs: 16 binary-classification runs (thumb vs. pinky) and 16 ternary-classification runs (thumb vs. index vs. pinky). To enable within-session adaptation, a base model was used to decode the first 8 runs of each task, after which a subject-specific fine-tuned model, updated using same-day data, was used for the remaining 8 runs.

**Table 1.** Stroke patients’ clinical characteristics.

| **Subject ID** | **Age** | **Gender** | **Dominant Hand** | **Affected Hand** | **Time of Stroke** | **Lesion Information** | **Stroke Type** |
| --- | --- | --- | --- | --- | --- | --- | --- |
| SS01 | 68 | F | R | L | 2021 | Right frontal lobe;<br>right parietal lobe;<br>right occipital lobe | hemorrhagic |
| SS02 | 72 | M | R | L | 2021 | Right ventral pons | hemorrhagic |
| SS03 | 57 | M | R | R | 2018 | Left basal ganglia | ischemic |
| SS04 | 66 | F | R | R | 2016 | Left corona radiata | ischemic |
| SS05 | 83 | M | R | R | 2024 | Left dorsal medulla | ischemic |
| SS06 | 60 | F | R | N | 2015 | Left frontoparietal cortex | hemorrhagic |
| SS07 | 77 | M | R | N | 2010 | None | Transient Ischemic Attack (TIA) |
| SS08 | 65 | M | L | N | 2019 | Right occipital lobe | ischemic (with petechial hemorrhage) |
| SS09 | 72 | M | R | L | 2015 | Right lateral pons | hemorrhagic |

Real-time finger decoding performance was evaluated using majority-vote accuracy, with performance trends examined across sessions and model types. To further characterize post-stroke neural reorganization, we compared task-related electrophysiological activity, oscillatory dynamics, and functional connectivity between stroke participants and a healthy control cohort (n = 16) recorded under an identical experimental protocol, at both sensor and source levels [28]. The healthy control dataset is publicly available at: https://doi.org/10.1184/R1/29104040. For consistency with the stroke cohort, only control participants who completed five sessions were included in the present analysis.

### Real-time robotic finger control in naïve stroke survivors

Across all participants (n = 9), the group-averaged online finger motor imagery (MI) control accuracy reached 83.54% for the two-finger task and 61.43% for the three-finger task by the end of training (Fig. 1A). In the two-finger condition, decoding between the thumb and pinky exhibited relatively balanced performance, as reflected by comparable precision and recall values across classes (S1 Fig. A, C). In contrast, performance in the three-finger condition was more heterogeneous. The index finger showed the lowest recall and was most frequently misclassified as the thumb (S1 Fig. B, D), indicating increased class-specific confusion with higher task complexity. A two-way repeated measures Analysis of Variance (ANOVA) on the first five sessions’ results revealed a significant improvement in MI performance over sessions for both the binary (*F* = 10.299, *p* < 0.001) and ternary (*F* = 4.287, *p* = 0.003) paradigms, demonstrating progressive enhancement of control proficiency with training. Although session effects were robust, the main effect of model type did not reach statistical significance for either the binary (*F* = 3.058, *p* = 0.084) or ternary (*F* = 0.300, *p* = 0.585) tasks, suggesting comparable performance between the Base model and the Fine-tuned model.

**Fig. 1.**
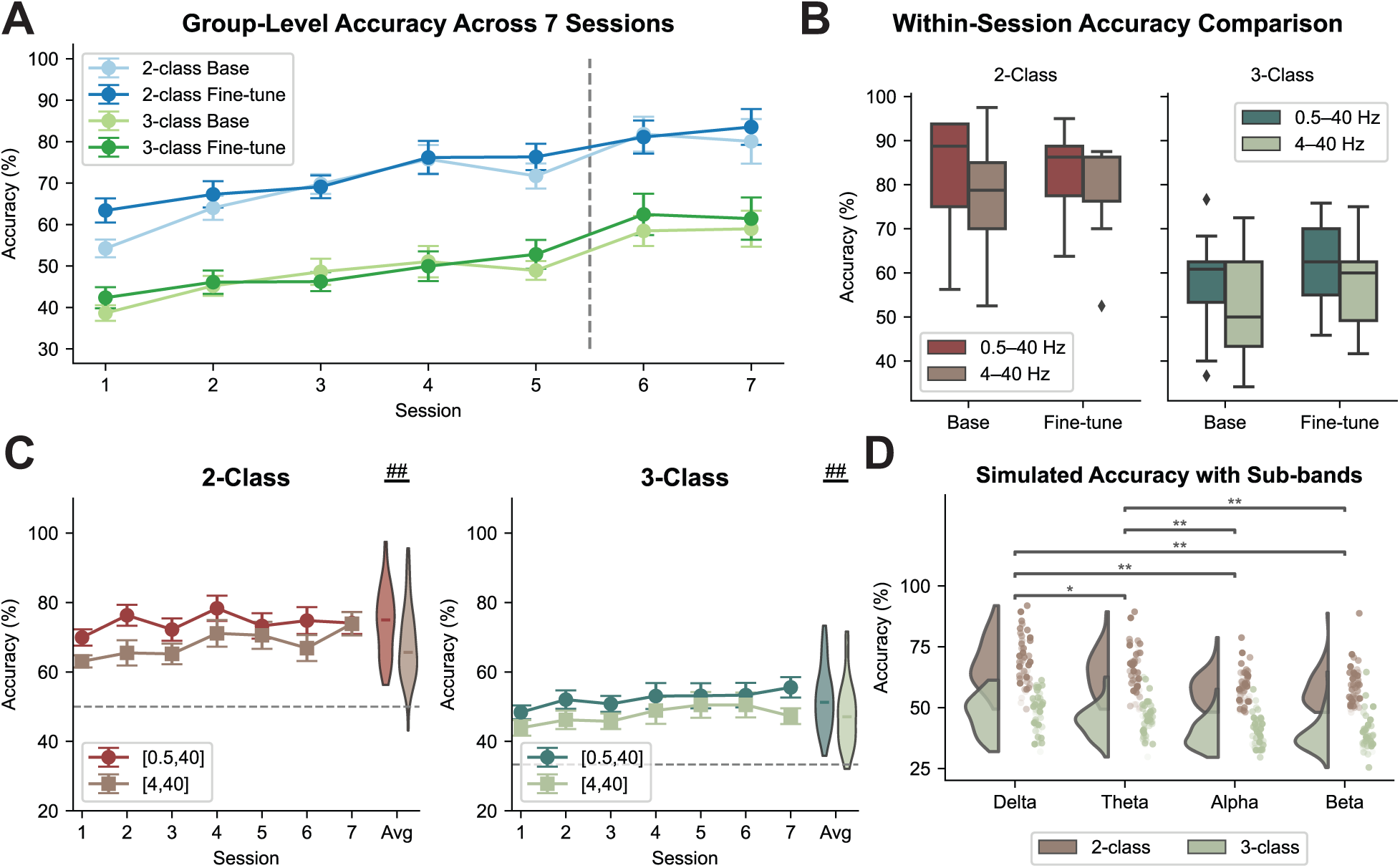
Online motor imagery (MI)–based robotic finger control performance and offline analysis of low-frequency contributions. **(A)** Group-level online performance for 2-finger and 3-finger MI across sessions (*n* = 9). The vertical gray dashed line represents the point when the online bandpass filtering parameter changed, and error bars indicate the standard errors. **(B)** Within-session online decoding accuracy comparing EEG inputs filtered at 0.5–40 Hz and 4–40 Hz (*n* = 9). The center lines indicate the median value. The boxes extend from the lower quartile to the upper quartile, and the lines indicate 1.5 times the interquartile range. Diamonds indicate outliers that are more than 1.5 times the interquartile range above the third quartile or below the first quartile. **(C)** Simulated offline fine-tuned decoding accuracy using EEG inputs filtered at 0.5–40 Hz and 4–40 Hz for 2-finger and 3-finger MI across seven online sessions (*n* = 9). The grey dashed line represents the chance level, and error bars indicate the standard error. The performance distribution across all sessions is shown on the right. The center lines indicate the median value. The boxes extend from the lower quartile to the upper quartile, and the lines indicate 1.5 times the interquartile range. Statistical analysis was performed using a two-way repeated-measures ANOVA with main effects of session and bandpass filtering setting. Significant main effects of bandpass are indicated (## if *p* < 0.01). **(D)** Simulated decoding accuracy using EEG signals filtered into individual frequency bands for the stroke survivors (delta, theta, alpha, and beta) (*n* = 9). Statistical significance was observed for the main effect of frequency bands in a two-way ANOVA. Significance stars indicate post hoc pairwise comparison results using an FDR-corrected two-tailed Wilcoxon signed-rank test (** if *p* < 0.01, * if *p* < 0.05).

Participants who performed MI control using their affected hand (n = 6) achieved average accuracies across all seven training sessions that were comparable to those of the non-affected hand group (n = 3), as reflected by low to moderate Cohen’s *d* values (Cohen’s *d* = 0.18 for 2-class; Cohen’s *d* = 0.42 for 3-class; S2 Fig. A). In contrast, participants exhibited superior decoding performance with their dominant hand (n = 6) relative to the non-dominant hand (n = 3), with moderate to large effect sizes (Cohen’s *d* = 0.78 for 2-class; Cohen’s *d* = 0.53 for 3-class; S2 Fig. B). Furthermore, a positive correlation was observed between arm-level MI performance, quantified by cursor-control task accuracy, and finger-level MI decoding performance (Pearson *r* = 0.636, S2 Fig. C, D), suggesting that proficiency in gross motor imagery generalizes to finer motor control representations.

### Low-frequency contributions to real-time finger MI decoding in individuals with stroke

A marked increase in online decoding performance was observed following the inclusion of low-frequency EEG components (0.5–4 Hz) in the decoding pipeline (Fig. 1A). To quantify the contribution of the delta band while accounting for learning-related effects across sessions, linear mixed-effects models were applied separately to each task condition, modeling changes in bandpass filtering as a fixed effect and session as a covariate. These analyses revealed a significant influence of the bandpass filtering setting on performance trends for both the three-class base model (*p* = 0.021) and the three-class fine-tuned model (*p* < 0.001), indicating that the inclusion of low-frequency components was associated with improved decoding performance beyond the potential learning effects involved in multi-session training. Numerical improvement in the decoding performance for the two-finger tasks was observed, but the effect of the bandpass filtering setting didn’t lead to statistical significance.

To further isolate the effect of low-frequency inputs, all nine participants completed an additional experimental session in which bandpass filtering settings (0.5–40 Hz vs. 4–40 Hz) were interleaved in a randomized order. The linear mixed-effects model revealed a significant main effect of bandpass (*z* = 2.124, *p* = 0.034, Fig. 1B) for the 3-class condition, indicating higher decoding accuracy when low-frequency components were included, providing within-session evidence supporting the contribution of low-frequency signals.

Consistent with the online results, offline simulations demonstrated significantly higher decoding accuracy for fine-tuned models trained on EEG signals filtered between 0.5–40 Hz compared with the 4–40 Hz condition for both two-finger and three-finger tasks, with similar trends observed for the base model (Fig. 1C and S3 Fig. A). A three-way repeated-measures ANOVA revealed a significant main effect of input frequency band on decoding accuracy (*F* = 22.453, *p* = 0.001). When decoding performance was evaluated using individual frequency bands, the delta band (0.5–4 Hz) contributed most strongly to decoding accuracy in the stroke group, followed by the theta band (4–8 Hz), whereas the alpha (8-13 Hz) and beta (13-30 Hz) bands contributed the least (*F* = 28.721, *p* < 0.001, Fig. 1D). This pattern differed from that observed in healthy participants (S3 Fig. B), in whom decoding performance was more strongly associated with alpha and beta band activity during continuous motor imagery tasks. Together, these results indicate a shift toward lower-frequency contributions to finger MI decoding after stroke, consistent with previously reported post-stroke shifts toward lower-frequency neural activity [30–32].

### Task-induced electrophysiological responses in stroke survivors compared with healthy controls

To characterize how stroke alters the cortical dynamics underlying MI, we examined both event-related desynchronization (ERD) in the alpha (8–13 Hz) and beta (13–30 Hz) bands, as well as movement-related cortical potentials (MRCPs) in the low-frequency range (0.3–3 Hz) during the finger MI task. Results of the stroke survivors were compared to those of 16 healthy controls originally collected from another study (*27*).

In stroke survivors, MI-induced suppression of oscillatory power exhibited an altered spatial distribution. Rather than the focal contralateral ERD typically observed in healthy participants, stroke survivors showed a more bilateral pattern of desynchronization, with increased involvement of the contralesional hemisphere (Fig. 2A, B). This spatial reorganization suggests greater engagement of the unaffected hemisphere during MI following stroke, potentially reflecting compensatory recruitment of preserved motor networks.

**Fig. 2.**
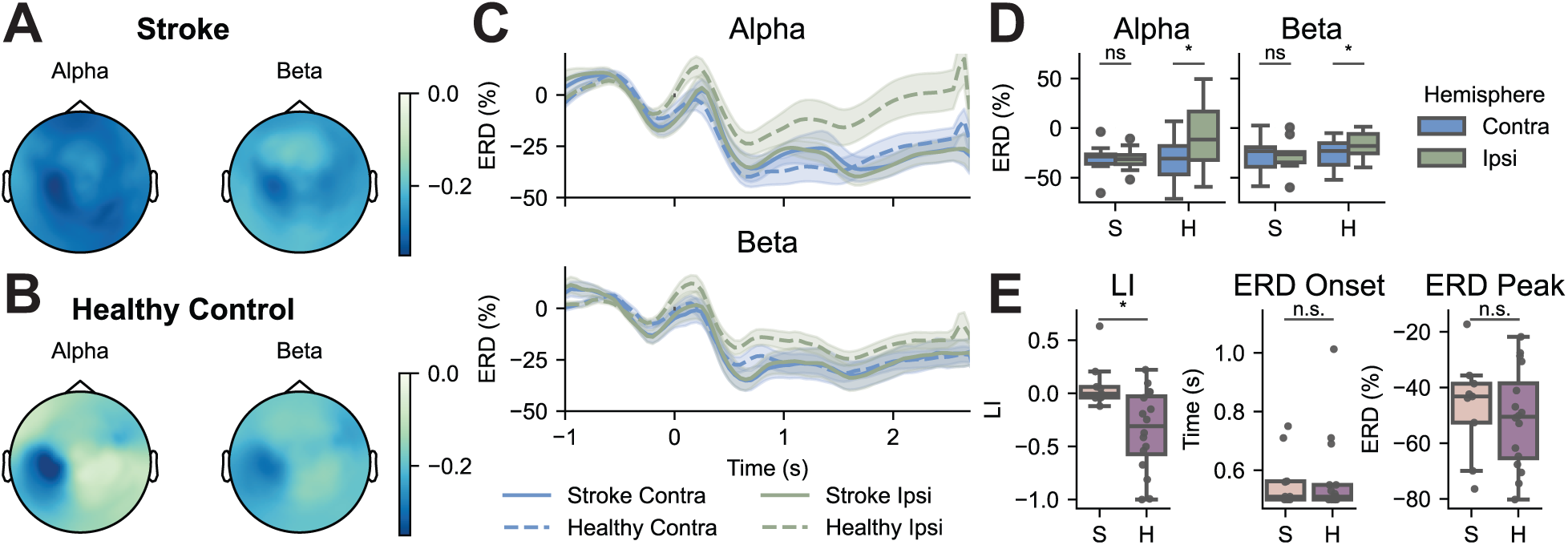
Comparison of movement-related cortical potential (MRCP) patterns between stroke survivors and healthy controls. (**A** and **B**) Group-averaged MRCP waveforms recorded over the contralateral sensorimotor cortex for different finger movements in healthy controls (**A**, *n* = 16) and over the ipsilesional sensorimotor cortex in stroke survivors (**B**, *n* = 9). The shaded area highlights a significant amplitude difference among the four fingers (p < 0.05, uncorrected for multiple comparisons), based on a one-way repeated-measures ANOVA. (**C** and **D**) Between-group comparisons of MRCP peak amplitude (**C**) and peak latency (**D**). Statistical differences were assessed using two-tailed Mann–Whitney U tests (** if *p* < 0.01).

Statistical analysis confirmed that, unlike the healthy group, where the contralateral ERD was significantly stronger than the ipsilateral (*p* = 0.018 for alpha; *p* = 0.026 for beta, Bonferroni-corrected), stroke participants showed no significant hemispheric difference (Fig. 2C, D). When comparing the laterality index, which quantifies the hemispherical ERD asymmetry, in stroke survivors with that in healthy controls, a significant difference was observed (*p* = 0.011, Fig. 2E). This loss of ERD laterality highlights a disruption in the normal contralateral-dominant sensorimotor activation pattern. In contrast, no significant group differences were observed in ERD onset or amplitude (Fig. 2E). Although ERD amplitude in stroke survivors was numerically lower than in controls, this reduction did not reach statistical significance.

Motivated by these differences in ERD topography between the stroke and healthy groups, we next examined the influence of regional EEG activity on decoding performance by restricting model inputs to specific cortical subregions. A near-significant main effect of cortical subregion on decoding accuracy was observed (*F* = 2.634, *p* = 0.072). However, among the four regions examined, EEG signals from the contralateral hand area did not produce the highest decoding performance (S4 Fig. A). This finding contrasts with the dominant contribution of contralateral motor regions observed in healthy individuals, suggesting increased recruitment of distributed and intact cortical areas during motor processing as a potential compensatory mechanism following stroke.

Notably, despite the contribution of low-frequency signals to decoding performance, no robust or spatially consistent ERD patterns were observed in the delta (0.5–4 Hz) or theta (4–8 Hz) frequency bands in either the stroke or healthy groups (S4 Fig. B, C).

Beyond alpha–beta oscillations, we further examined motor-related cortical potentials (MRCPs), which capture slow-cortical potentials associated with movement planning and initiation [33]. The low-frequency activity (0.3–3 Hz) during individual finger MI was extracted from offline sessions and compared between groups. In healthy controls, MRCPs were clearly observable as a negative deflection occurring shortly after trial onset, with finger-specific variations evident within a brief post-onset window (Fig. 3A). In contrast, stroke participants exhibited greater inter-individual variability and lacked a consistent negative deflection at the group level (Fig. 3B, C). The peak MRCP amplitude was significantly attenuated, and the peak latency was significantly prolonged relative to the control group, suggesting delayed cortical engagement and impaired neural responsiveness. (Fig. 3C, D).

**Fig. 3.**
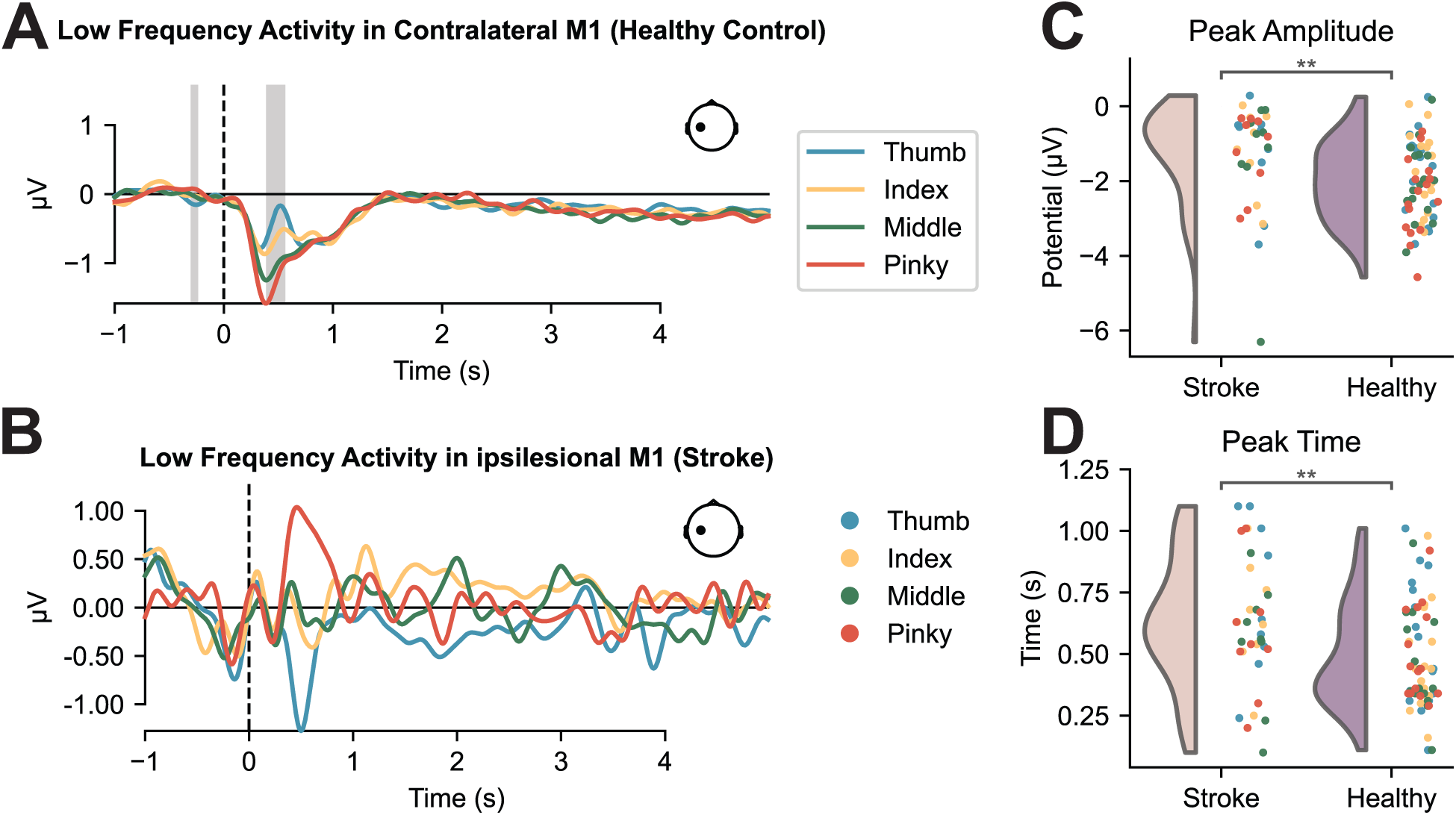
Comparison of event-related desynchronization (ERD) patterns between stroke survivors and healthy controls. (**A** and **B**) Group-averaged topographical maps of alpha-band (8–13 Hz) and beta-band (13–30 Hz) ERD during online MI control for the stroke group (**A**) and the healthy control group (**B**). (**C**) Temporal evolution of alpha- and beta-band ERD, averaged across channels of interest (contralateral and ipsilateral sensorimotor regions relative to the control hand). Solid lines indicate group means, and shaded areas denote the standard error of the mean. (**D**) Group-averaged ERD magnitude in the contralateral and ipsilateral sensorimotor regions relative to the control hand. The center lines indicate the median value. The boxes extend from the lower quartile to the upper quartile, and the lines indicate 1.5 times the interquartile range. Circles indicate outliers that are more than 1.5 times the interquartile range above the third quartile or below the first quartile. Hemispheric differences were assessed using two-tailed Wilcoxon signed-rank tests with Bonferroni correction for multiple comparisons (* if *p* < 0.05, n.s. if no statistical significance is found). (**E**) Laterality index (LI), ERD onset time, and peak contralateral ERD magnitude. Between-group comparisons were performed using two-tailed Mann–Whitney U tests (* if *p* < 0.05, n.s. if no statistical significance is found).

Together, these findings demonstrate that stroke alters both the oscillatory and low-frequency electrophysiological correlates of MI. The combination of attenuated ipsilesional ERD, weakened and delayed MRCPs, and enhanced contralesional recruitment reflects a complex interplay between functional impairment and adaptive neuroplasticity. The bilateral activation and reduced MRCP amplitude suggest that sensorimotor network reorganization after stroke may involve a redistribution of cortical resources, where the contralesional motor cortex compensates for impaired ipsilesional pathways.

### Altered cross-frequency dynamics and functional connectivity following stroke

To further investigate network disruption and reorganization in stroke survivors, we examined cross-frequency interactions focusing on the alpha and beta bands. Expanding from local coupling, we next assessed inter-regional functional connectivity and global spectral patterns in source-level EEG to characterize stroke-related changes at the large-scale network level.

A strong positive correlation was found between contralateral alpha ERD and beta ERD in healthy controls (Pearson *r* = 0.89, *p* < 0.001), whereas this relationship was weakened in the stroke group (Pearson *r* = 0.75, *p* = 0.019; Fig. 4A). These results indicate an alternated coordination between alpha and beta desynchronization, suggesting modified coupling within the sensorimotor rhythm following stroke.

**Fig. 4.**
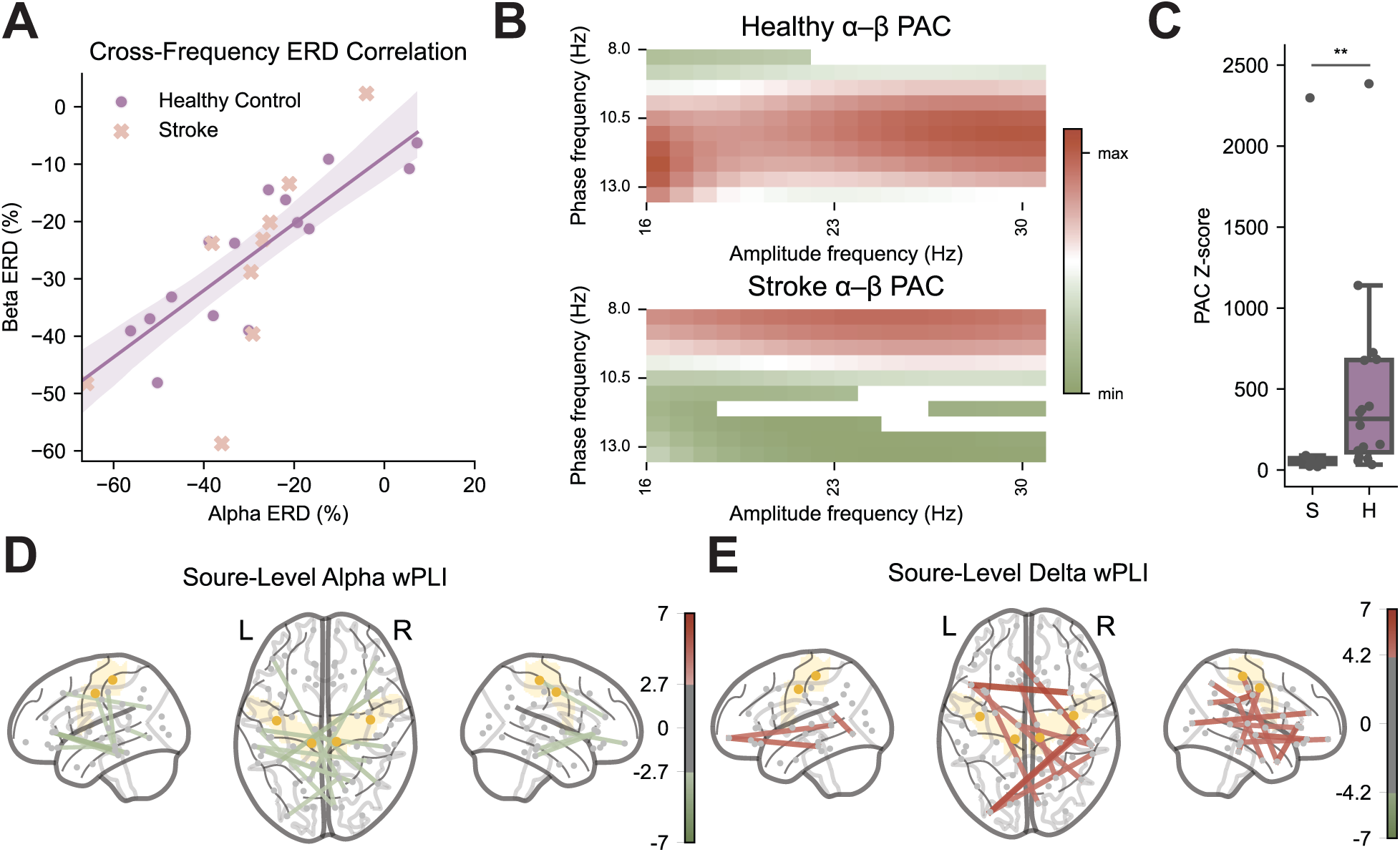
Altered cross-frequency dynamics and functional connectivity in stroke survivors. (**A**) Cross-frequency coupling between alpha- and beta-band ERD over the contralateral sensorimotor cortex in healthy controls and over the ipsilesional sensorimotor cortex in stroke survivors. The solid line represents the linear regression fit to the healthy control data, and the shaded region denotes the 95% confidence interval. (**B**) The alpha-beta phase-amplitude coupling (PAC) of the stroke and healthy group over the contralateral sensorimotor cortex (the ipsilesional sensorimotor cortex for stroke survivors). PAC values were normalized to the maximum value observed across the two groups for visualization. (**C**) Comparison of PAC z-scores between groups. Z-scores were computed based on surrogate distributions generated using 200 permutations. Between-group differences were assessed using two-tailed Mann–Whitney U tests (** if *p* < 0.01). (**D** and **E**) Group-level differences in alpha-band (**D**) and delta-band (**E**) weighted Phase Lag Index (wPLI) at the source level, expressed as z-scores relative to healthy controls. Functional connections between the atlas with absolute z-scores exceeding the 99th percentile threshold are displayed, with edge color and thickness indicating the sign and magnitude of the z-score. L and R refer to the contralateral side (the ipsilesional side for stroke survivors) and the ipsilateral side (the contralesional side for stroke survivors) respectively.

Consistent with this observation, phase–amplitude coupling (PAC) between the alpha and beta bands in the contralateral M1 region (corresponding to the ipsilesional side in stroke) was significantly reduced in stroke survivors (Fig. 4B, C), further confirming impaired cross-frequency communication within the motor cortex. Interestingly, we observed that in the healthy control group, beta activity is mainly correlated with high alpha oscillations, while in the stroke group, a lower frequency range, including low alpha and theta, shows higher correlation with the beta activities, which might suggest cortical slowing after stroke (Fig. 4B, S5 Fig.).

To further characterize stroke-related changes in neural activity across both spatial and spectral domains, we performed source localization to project EEG signals from the sensor level onto the cortical surface and computed functional connectivity within individual frequency bands. At the network level, a widespread reduction in alpha-band functional connectivity and a compensatory increase in delta-band connectivity were observed after stroke (Fig. 4D, E). In particular, alpha connectivity was markedly reduced within the ipsilesional hemisphere and across hemispheres, whereas increased delta connectivity was primarily observed in the contralesional hemisphere, possibly reflecting adaptive recruitment of intact motor circuits. Together, these results highlight a disrupted alpha-mediated integration and strengthened delta synchronization, indicating both network disintegration and compensatory reorganization following stroke. This reorganization also provides electrophysiological support for the observed shift in decoding contributions from the alpha to the delta band. Similarly, functional connectivity in the theta and beta bands exhibited contralesional enhancement patterns (S6 Fig.).

Previous studies have consistently reported an increased delta–alpha ratio (DAR) following stroke, reflecting the characteristic slowing of cortical rhythms after ischemic injury [34]. Elevated delta activity has been associated with neuronal deafferentation, whereas reduced alpha power indicates impaired large-scale network integration [31,35]. To examine these alterations in our cohort, we compared delta power, alpha power, and DAR between stroke survivors and healthy controls. A globally elevated DAR was observed in the stroke group, with a more pronounced increase localized to the ipsilateral sensorimotor cortex (corresponding to the contralesional hemisphere for stroke patients; Cohen’s *d* = 0.94, Fig. 5A, B). Compared with controls, delta power was higher on the contralateral (ipsilesional) side and more symmetrically distributed across hemispheres in stroke survivors (Fig. 5C, D), while alpha power showed a selective decrease over the ipsilateral (contralesional) side but remained relatively preserved over the contralateral (ipsilesional) hemisphere (Fig. 5E, F). This alpha pattern deviates from prior resting-state findings, where alpha suppression typically occurs on the lesioned side [36,37]. This discrepancy likely arises from the task context. DAR here was computed during motor imagery rather than rest, such that the healthy group’s contralateral alpha ERD may have lowered their baseline reference. Consequently, the observed pattern suggests both a reduced baseline alpha power and a diminished capacity for alpha modulation in stroke patients during motor imagery.

**Fig. 5.**
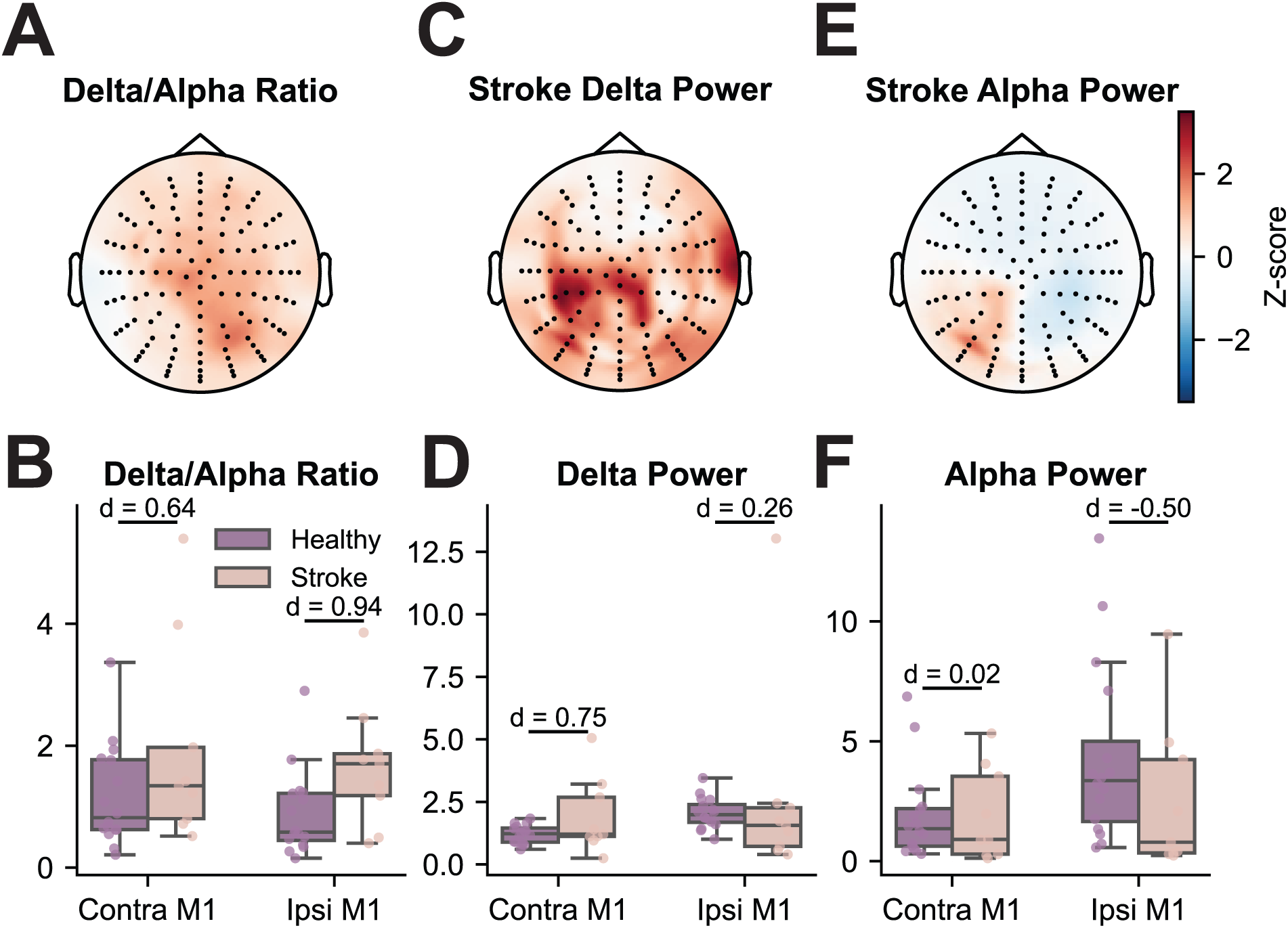
Alterations in delta–alpha power balance following stroke. **(A–C)** Group-averaged topographical maps of the delta–alpha ratio (DAR; **A**), delta-band power (**B**), and alpha-band power (**C**) in stroke survivors, expressed as z-scores relative to the healthy control baseline. **(D–F)** Between-group comparisons of DAR (**D**), delta power (**E**), and alpha power (**F**) over the contralateral and ipsilateral sensorimotor cortices. Differences across groups are quantified using Cohen’s *d*.

## Discussion

This study extends recent development of EEG-based robotic finger control to BCI-naïve stroke populations, showing that fine-grained motor imagery (MI)–based control can be achieved without prior exposure to similar paradigms. Averaged online decoding performances reached 83.54% for two-finger control and 61.43% for three-finger control, which were comparable to those obtained from the BCI-experienced healthy cohort. These results suggest that the proposed deep-learning-based finger decoding paradigm may generalize beyond expert users and is robust to inter-individual variability and neurological impairment. Importantly, the ability of stroke survivors to achieve reliable control by imagining movements of the impaired hand, despite structural brain injury and motor deficits, provides a critical foundation for translating this system toward assistive robotic applications aimed at restoring functional hand use in daily life, consistent with prior demonstrations of MI-based arm-level BCI control in post-stroke populations [16,19].

Despite the comparable decoding performance, the underlying neural signatures differed significantly between cohorts. Comparisons of task-elicited neural responses between stroke survivors and healthy participants revealed clear alterations in the spatiotemporal characteristics of MI–related cortical dynamics. Specifically, stroke survivors exhibited a more bilateral pattern of ERD, accompanied by a reduced amplitude of the ERD response. In addition, MRCP responses were attenuated and delayed relative to healthy controls. Increased bilateral recruitment of sensorimotor areas during movements of the affected hand after stroke has been consistently reported across neuroimaging studies, and is commonly interpreted as a compensatory response to reduced efficiency of the ipsilesional motor system [38,39]. Such bilateral activation patterns contrast with the focal contralateral ERD typically observed in healthy individuals and reflect stroke-induced reorganization of motor networks.

The reduced amplitude of ERD observed in stroke survivors aligns with prior work showing attenuated movement-related beta and alpha desynchronization in patients post-stroke compared with healthy controls, with lower ERD amplitudes correlating with the degree of motor impairment [40,41]. Attenuated ERD has been associated with a decreased capacity to suppress ongoing oscillatory activity during motor preparation and imagery, potentially reflecting decreased GABA-A receptors or GABA concentration affected by the lesions. The delayed MRCP onset observed in the stroke group further aligns with reports of slowed cortical dynamics and disrupted timing of sensorimotor processing after stroke [40,42]. Additionally, greater inter-individual variability was observed in the MRCP responses in the stroke group, which might be due to the heterogeneous nature of stroke lesions [43].

Stroke can induce widespread neural reorganization beyond the lesioned motor region, including contralesional and brain-wide synaptic remodeling [44]. At the network level, our results showing a widespread reduction in alpha-band functional connectivity after stroke are consistent with previous electrophysiological and neuroimaging studies [36,39]. Reduced functional connectivity within M1 has been frequently reported as a function of motor impairment, suggesting compromised coordination between motor cortices after stroke, therefore contributing to impaired sensorimotor integration and motor deficits [45–47]. Conversely, the enhanced delta-band connectivity, especially within the contralesional hemisphere, may reflect compensatory recruitment of alternative motor circuits when ipsilesional pathways are compromised due to structural brain injury [46,48]. This pattern of reduced alpha connectivity and heightened delta synchronization suggests both network disintegration and compensatory reconfiguration after stroke.

The elevated delta–alpha ratio observed in stroke survivors compared with healthy controls is consistent with the literature linking increased DAR to post-stroke cortical dysfunction. DAR has been used as a quantitative EEG marker to discriminate between ischemic stroke patients and controls, with elevated delta and reduced alpha power, reflecting the spectral slowing [34,49]. Structural damage in the lesioned area, causing disruption of thalamocortical connections, has been associated with increased power in the low frequency band and suppression of fast activity [50,51]. Increased delta activity after stroke has been principally associated with tissue dysfunction and related impairment, which is consistent with our finding that ipsilesional delta power was higher and more symmetrically distributed across hemispheres in stroke survivors [49]. The reduction of alpha power in stroke patients that was only observed in the contralesional side of the brain likely reflects the task-specific nature of motor imagery. In our control group, pronounced contralateral alpha ERD likely lowered the alpha power estimates, revealing a relatively preserved alpha power that contrasts with typical resting-state findings where alpha suppression is often most prominent over the lesioned hemisphere [52,53].

Interestingly, a key finding of this study is the pivotal role of low-frequency oscillations in decoding finger movements for stroke survivors, a pattern that was not observed in the healthy population, in which alpha and beta oscillations contributed most strongly to decoding performance. Although previous research has highlighted the importance of low-frequency components for movement decoding in general [54–56], the prominent contribution of these low-frequency components in our stroke cohort likely reflects stroke-specific electrophysiological alterations rather than general MI phenomena. Literature has characterized a spectral slowing phenomenon in stroke, whereby the power spectrum shifts toward lower frequencies and elevated low-frequency power is detected, reflecting both periodic and aperiodic neural abnormalities in stroke populations [49]. In line with this framework, our findings, including increased delta-band power, enhanced delta-band functional connectivity, and reduced alpha-band integration, suggest that motor-related information in stroke is increasingly encoded within slower oscillatory dynamics. These results provide electrophysiological evidence that the observed shift in decoding contributions from alpha to delta bands reflects an underlying reorganization of neural dynamics.

Despite the altered spectral signatures, decoding performance remained robust, underscoring the ability of deep learning–based models to extract discriminative information from distributed and reorganized neural representations. This dissociation between altered electrophysiological signatures, including the reduced contralateral ERD and attenuated movement-related cortical potentials, and decoding performance underscores the distinction between neural efficiency and decodability. Indeed, prior MI-BCI studies have shown that deep learning decoders exploit the EEG task-relevant features across broader spatial distributions and different frequency bands beyond typical sensorimotor rhythmic features, particularly when adaptive classifiers or data-driven feature learning approaches are employed to accommodate heterogeneous neural patterns [28,57].

More broadly, our findings support that stroke-related neural reorganization, although it results in more heterogeneous EEG signals compared to healthy controls, can nonetheless provide a viable substrate for BCI control. The present results suggest that such compensatory dynamics are effectively leveraged by flexible, data-driven models. Deep learning frameworks not only enable the automatic extraction of dynamic and non-linear EEG features that are difficult to capture with traditional decoding pipelines, but also support individualized and adaptive decoding architectures that are well-suited to heterogeneous and reorganized neural activity patterns in stroke, thereby holding substantial promise for real-world BCI-based clinical applications [58].

The present findings can also be interpreted within the broader context of noninvasive BCI applications for motor control. Prior studies have established the feasibility of controlling external devices, including robotic arms [59,60], wheelchairs [61], and drones [62], primarily using control signals derived from gross motor imagery. In contrast, finger-level decoding demands substantially greater control granularity. Despite this increased complexity, we demonstrate that stroke survivors can achieve comparable decoding performance without prior BCI experience, supporting the feasibility of higher-dimensional neural control and advancing toward more dexterous robotic assistive technologies.

Overall, these decoding analyses indicate that both spectral composition and regional signal selection influence finger-specific MI decoding performance in the stroke group. The observed differences in frequency-band and regional contributions, relative to healthy controls, suggest that stroke survivors engage motor imagery through reorganized and distributed cortical networks, which is consistent with broader evidence of motor network plasticity after stroke. However, several limitations require consideration. One limitation of the study is the age difference between the stroke group and the healthy control group. Even though the user’s age and BCI performance are found to be not significantly correlated, this may still confound interpretations of neurophysiological differences and requires careful consideration in future work [63,64]. In addition, the optimization of parameters during the online experiments may have influenced learning curves. Future studies utilizing fixed, optimized parameters in larger, naïve populations are needed to precisely quantify the training responses. Furthermore, a subset of stroke participants did not exhibit overt motor impairment, which may introduce additional heterogeneity in the cohort. At the same time, this reflects the clinical variability of stroke and provides an opportunity to examine whether BCI performance is sensitive to subclinical neurophysiological alterations beyond observable motor deficits. In the present study, no significant difference in BCI performance was observed between participants with and without motor impairment. While this finding suggests that the decoding approach may be applicable across a range of functional states, further validation in larger and more stratified cohorts is needed. Overall, these results indicate the potential of deep learning -based methods to bridge the gap between disease-affected brain electrical activity and sophisticated functional robotic control across heterogeneous neurophysiological conditions.

## Materials and Methods

### Experimental design

#### Subjects

Ten stroke-affected individuals were enrolled in this study under a protocol approved by the Carnegie Mellon University Institutional Review Board (STUDY2017_00000548). All participants completed eligibility screening and provided written informed consent. One participant withdrew due to scheduling conflicts; nine completed the full study (6 male / 3 female, age: 68.9 ± 8.1 years; eight right-handed). Each participant attended two sessions of limb-level MI cursor training, followed by two finger-MI offline sessions and five finger-MI online sessions with robotic feedback.

#### EEG acquisition and setup

EEG was recorded using a 128-channel BioSemi ActiveTwo system (BioSemi, Amsterdam, Netherlands) placed using the international 10–20 positioning scheme. Data were sampled at 1,024 Hz. Participants sat comfortably at a distance of ∼90 cm from a display, while an Allegro robotic hand (Wonik Robotics, Korea) was positioned between the subject and the monitor to provide visual feedback of the decoded finger movements. Subjects were instructed to perform kinesthetic motor imagery while avoiding overt movements.

#### Limb-level cursor MI training task

To establish baseline MI control proficiency, participants performed two sessions of 1-D cursor control (Left–Right and Up–Down). Each session consisted of 8 runs per task, with 25 trials in each run. Subjects are instructed to imagine moving their right hand to move the cursor to the right and their left hand to move it to the left. For upward movement, they imagine moving both hands, while for downward movement, they are instructed to relax. Each trial began with a visual cue indicating the target location, followed by real-time cursor movement feedback for up to 6 seconds. A surface Laplacian was applied to channels around C3 and C4, and high alpha-band power (10.5 – 13.5 Hz) was estimated using an autoregressive model. A linear classifier computed hemisphere-specific power differences (C4–C3 for LR, -C3-C4 for UD), and the normalized output controlled cursor direction and speed. If the cursor reaches the visible target within this time, the trial is considered as a hit. If the cursor moves in the opposite direction and reaches an invisible target, the trial is counted as a miss. If the cursor ended up somewhere in between the two targets without hitting either side, the trial is marked as aborted. Accuracy was defined as hits divided by the number of hit and miss trials, excluding aborted trials.

#### Finger motor imagery without feedback

After cursor training, participants completed two sessions of finger-specific MI without feedback. They imagined repetitive flexion and extension of one of four fingers (thumb/index/middle/pinky) of their dominant hand if they did not have motor impairments (n = 3) or their affected hand (n = 6). Each session consisted of 32 runs, each containing 5 trials for each finger in a randomized order. Trials lasted 5 s with 2 s inter-trial intervals.

#### Finger motor imagery with real-time robotic feedback

During the online sessions, participants performed finger MI tasks with real-time robotic feedback, following the paradigm introduced in Ding et al. [28]. Each session consisted of 32 runs, alternating between binary and ternary classification tasks that decoded imagined finger movements into robotic hand motions. In the ternary condition, subjects imagined flexion and extension of the thumb, index, or pinky finger, while in the binary condition, they imagined thumb and pinky movements. The first half of each session used a base model trained on data collected in previous sessions, while the second half employed a fine-tuned model that was updated using data from the same day’s first eight runs.

Each trial lasted three seconds. Participants initiated self-paced finger imagery at the trial onset. After one second, robotic feedback began and continued for the remaining two seconds. The Allegro robotic hand (Wonik Robotics, Korea) provided immediate visual feedback by flexing the finger predicted by the decoder, updating every 125 ms. Concurrently, a screen display indicated the decoder’s decision with the target finger turning green for correct classifications and red for incorrect ones. The finger that accumulated the greatest flexion by the end of the feedback period was considered as the final prediction for that trial.

Online processing and classification were performed using custom Python (3.8.4) scripts made for BCPy2000 (2021.1.0), a part of the BCI2000 program [65]. The 128-channel EEG data were re-referenced to the common average, downsampled to 100 Hz, and bandpass-filtered between 4 and 40 Hz for the first five sessions and between 0.5 and 40 Hz for the last two sessions. For each 125 ms update, the most recent one-second window was z-scored and fed into the decoder. Real-time classification was performed using EEGNet-8,2 with an online smoothing approach to enhance the stability of control outputs [28,29]. Five participants participated in an additional experimental session in which bandpass filtering settings (0.5–40 Hz vs. 4–40 Hz) were interleaved in a randomized order.

#### Task-induced electrophysiological analysis

EEG data during the online finger MI sessions were processed for event-related desynchronization (ERD) analysis to uncover the electrophysiological patterns associated with finger MI. This was done using the FieldTrip (20230118) toolbox [66] and customized MATLAB (R2023a) scripts (MathWorks Inc., MA, USA). The raw EEG data were re-referenced to the common average, downsampled to 100 Hz, and bandpass filtered between 0.5 and 30 Hz. Independent component analysis (ICA) was performed to remove eye movement and muscle artifacts. The continuous signal was then segmented into trials spanning from 2 seconds before trial onset to the end of the trial. To calculate ERD for each EEG channel, Morlet wavelets were used to extract the average power in the alpha (8 - 13 Hz) and beta (13 - 30 Hz) bands from 0.5 seconds after trial onset until the end of the trial. The average power within the same frequency bands during the 1-second period preceding trial onset was used as the baseline. Single-trial ERD was quantified by the relative change in alpha and beta band power, as calculated in Eq. 1.

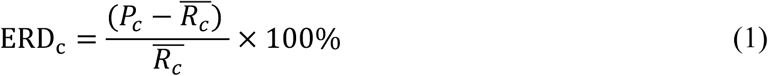

where *P*_c_ represents the average power in the alpha and beta bands during the task period, and *R*_c_ denotes the average baseline power within each session. The ERD activities in channel C3 and C4 were selected to represent either the contralateral or ipsilateral motor cortex activities, depending on the control hand. The metrics that were used for comparison between the stroke group and the healthy population include laterality index (LI), ERD onset, and ERD peak. LI is defined as the difference between the contralateral mean ERD and the ipsilateral mean ERD divided by the sum of the absolute values of the contralateral and ipsilateral ERD. ERD onset is defined as the first time that the ERD value exceeds -20% and is sustained for at least 0.1 second, while the minimum ERD value during the entire window is considered the ERD peak value.

Raw EEG data were further processed to visualize MRCPs in the low-frequency band (0.3–3 Hz). Data processing and visualization were conducted using MNE-Python (1.9.0) [67]. The EEG signals were re-referenced to the common average, downsampled to 100 Hz, and bandpass filtered between 0.3 and 3 Hz. ICA was performed to remove eye movement artifacts. The continuous data were segmented into trials, spanning from 1 second before trial onset to the end of each trial. The signal during the second before the trial onset was used for baseline correction. Group-averaged MRCPs within the contralateral motor region (either C3 or C4) were plotted for each finger movement. Time periods showing significant amplitude differences among the four fingers were determined using a one-way repeated-measures ANOVA. The peak MRCP amplitude and time relative to the trial onset were compared between groups through a two-way mixed model ANOVA.

#### Spectral dynamics analysis

The correlation between the contralateral alpha and beta ERD was evaluated and quantified through a linear regression. The regression residual is defined as the absolute deviation between the fitted value and the actual value.

Phase–amplitude coupling (PAC) reflects a form of nonlinear neural coupling in which the phase of a slower rhythm modulates the amplitude envelope of a faster oscillation [68]. PAC analyses were performed using the pactools Python toolbox [69]. EEG signals were first aligned across subjects by horizontally flipping channel indices for stroke-affected participants whose control hand was not the right hand, ensuring consistent hemispheric correspondence. For each subject, signals were averaged across electrodes corresponding to the contralateral M1 region of interest (either C3 or C4). PAC was computed using the Tort modulation index method [68]. Phase-providing frequencies were sampled within the delta–theta (0.5–8 Hz) and alpha (8–13 Hz) bands, while amplitude-providing frequencies were sampled within the beta band (13–30 Hz). For each phase–amplitude frequency pair, surrogate distributions were generated using 200 permutations to estimate statistical significance. Group-averaged Z-scored PAC values relative to the surrogate distribution were visualized. For visualization purposes, the amplitude frequency range was restricted to 16–30 Hz to reduce the influence of within-band or adjacent-band interactions.

To characterize alterations in the spectral activities after stroke, the delta and alpha power were computed from the FieldTrip preprocessed EEG data using a Welch-based spectral analysis. From these values, the delta–alpha ratio (DAR), which is defined by the delta power divided by alpha power, was derived. Differences in channel-wise power and DAR between the healthy and stroke groups were quantified using z-scores.

#### Source-level functional connectivity analysis

Source-level functional connectivity was computed from reconstructed source waveforms at regions of interest (ROIs) [70] with the aid of MNE-Python (1.9.0) [67]. Preprocessed EEG epochs were projected onto a standard BioSemi 128-channel montage and mapped onto a template anatomy based on the *fsaverage* brain. Forward solutions were computed using a three-layer boundary element model, and cortical source activity was reconstructed using the dynamic statistical parametric mapping (dSPM) inverse solution.

Cortical source time series were extracted from ROIs based on the Desikan–Killiany atlas [71]. To account for differences in lesion laterality and ensure comparability across subjects, left and right hemispheric ROIs were swapped when necessary such that all data were aligned relative to the contralateral hemisphere of the control hand.

Functional connectivity within individual frequency bands was quantified using the weighted Phase Lag Index (wPLI) approach, which estimates the consistency of non-zero phase-lag relationships between signals while reducing the influence of volume conduction and common reference effects [72,73]. EEG connectivity was computed using the MNE-connectivity toolbox [74]. Spectral connectivity was estimated using a Fourier-based approach, with wPLI computed separately within the delta (0.5–4 Hz), theta (4–8 Hz), alpha (8–13 Hz), and beta (13–30 Hz) bands. Within each band, connectivity values were averaged across frequencies, yielding one ROI-to-ROI connectivity matrix per band for each subject.

For group-level analysis, permutation-based z-score maps were computed to assess differences in wPLI between healthy controls and stroke participants. Only connections whose absolute z-scores exceeded the 99th percentile threshold were displayed on the cortical surface, with edge color reflecting the magnitude of the corresponding z-scores.

#### Online decoding simulation

Data collected during the online sessions were further evaluated offline using EEGNet-8,2 to assess the discriminability of individual finger motor imagery across different frequency components and cortical subregions. The offline preprocessing pipeline, model training, and evaluation procedures were identical to those used in the online decoding framework, enabling a direct simulation of online decoding performance. EEG signals were band-pass filtered using multiple frequency ranges, including 4–40 Hz, 0.5–40 Hz, delta (0.5–4 Hz), theta (4–8 Hz), alpha (8–13 Hz), and beta (13–30 Hz) bands. In addition, different subsets of EEG electrodes were selected as model inputs, including contralateral and ipsilateral hand-knob regions, parietal regions, and frontal regions, to examine region-specific cortical contributions on decoding performance.

### Statistical analysis

Statistical analyses were performed using custom Python (version 3.10.13) scripts. Unless otherwise specified, paired performance metrics and electrophysiological measures were compared using Wilcoxon signed-rank tests. Differences in behavioral and electrophysiological metrics between the healthy control and stroke groups were assessed using Mann–Whitney U tests. Bonferroni correction was applied to control for multiple comparisons where applicable.

Two-way repeated-measures analyses of variance (ANOVAs) were conducted to evaluate the main effects of session and condition (model type or band-pass filtering setting) on both online decoding performance and simulated online decoding results comparing the 0.5-40 Hz and the 4-40 Hz inputs. Three-way repeated-measures ANOVAs were used to assess the main effects of session, number of classes, and condition (band-pass filtering setting or cortical subregion) on simulated online decoding performance using individual frequency bands and region-specific electrode subsets. When a significant main effect was observed (*p* < 0.05), pairwise post hoc comparisons were performed using Wilcoxon signed-rank tests with false discovery rate (FDR) correction.

## Acknowledgments

We thank our former colleague, Dr. Dylan Forenzo, for setting up the IRB protocol recruiting stroke participants, and Chalisa Udompanyawit, for their contributions to setting up the robotic interface. We also gratefully acknowledge Pitt+Me for their support in recruiting stroke participants.

This work was supported in part by National Institutes of Health grants NS124564 (BH), NS131069 (BH), NS127849 (BH), and NS096761(BH). Y.D. was supported in part by a Carnegie Mellon University’s Center for Machine Learning and Health Fellowship. M.K. was partially supported by National Institutes of Health T32 training grant EB029365 (PI: BH).

## Author contributions

Conceptualization: YD, BH. Methodology: YD, MK, HW, BH. Investigation: YD, MK, ZJ, HW, JZ, BH. Formal analysis: YD. Visualization: YD. Resources: GW, BH. Supervision: BH. Writing—original draft: YD. Writing—review & editing: YD, GW, BH.

## Competing interests

Authors declare that they have no competing interests.

## Data and materials availability

All data are available in the main text or the supplementary materials. Additional EEG data in all subjects will be made available in a public online repository when the paper is published. Customized computer codes will be made public on GitHub when the paper is published.

## Supporting Information

**S1 Fig.**
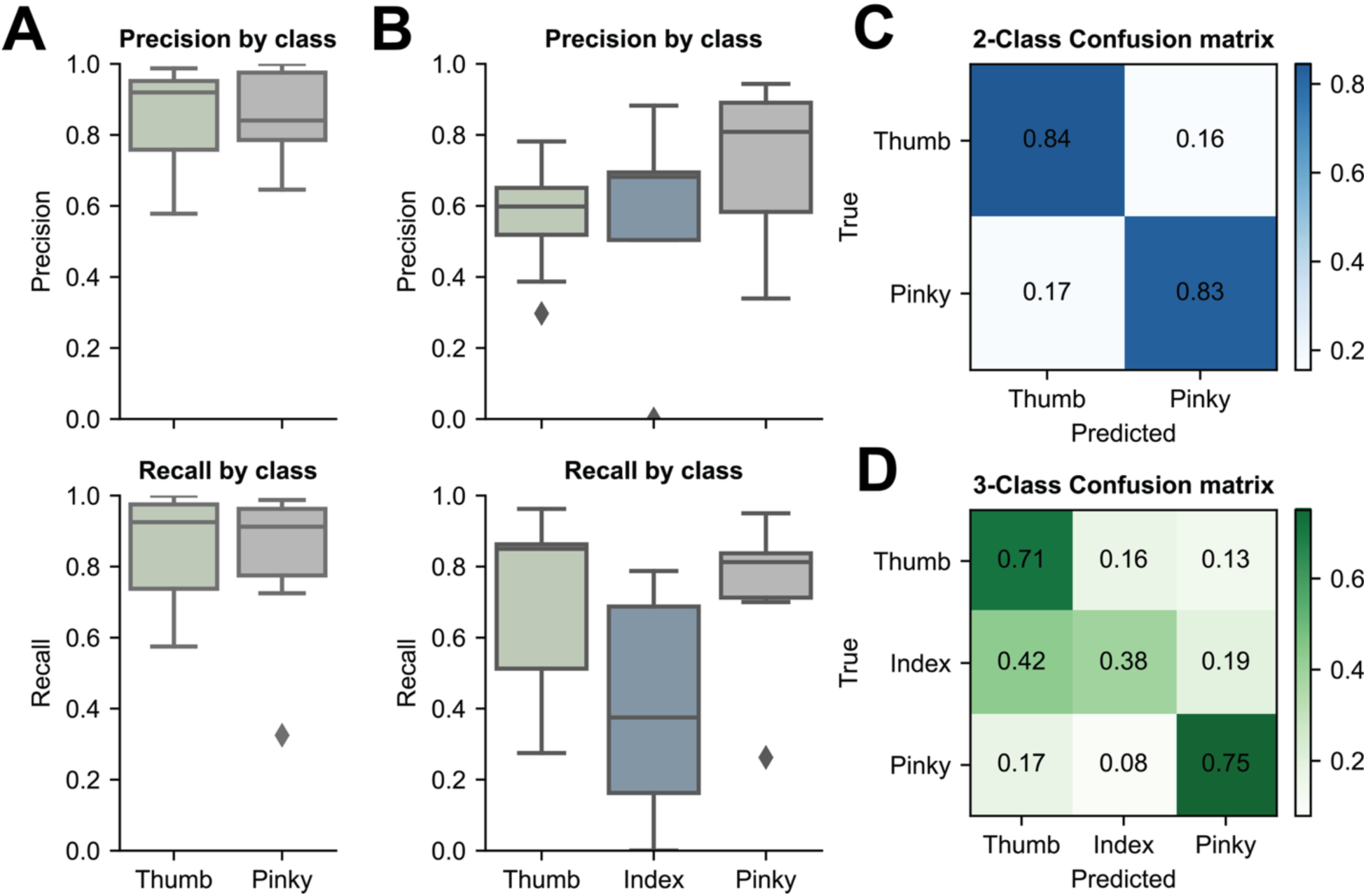
Online motor imagery (MI)–based robotic finger control performance (*n* = 9). (**A** and **B**) Group-level precision and recall of 2-finger (**A**) and 3-finger (**B**) decoding. The center lines indicate the median value. The boxes extend from the lower quartile to the upper quartile, and the lines indicate 1.5 times the interquartile range. Diamonds indicate outliers that are more than 1.5 times the interquartile range above the third quartile or below the first quartile. (**C** and **D**) Confusion matrices for 2-finger (**C**) and 3-finger (**D**) online decoding results. The values represent the proportion of predictions for each true label classified as each predicted label.

**S2 Fig.**
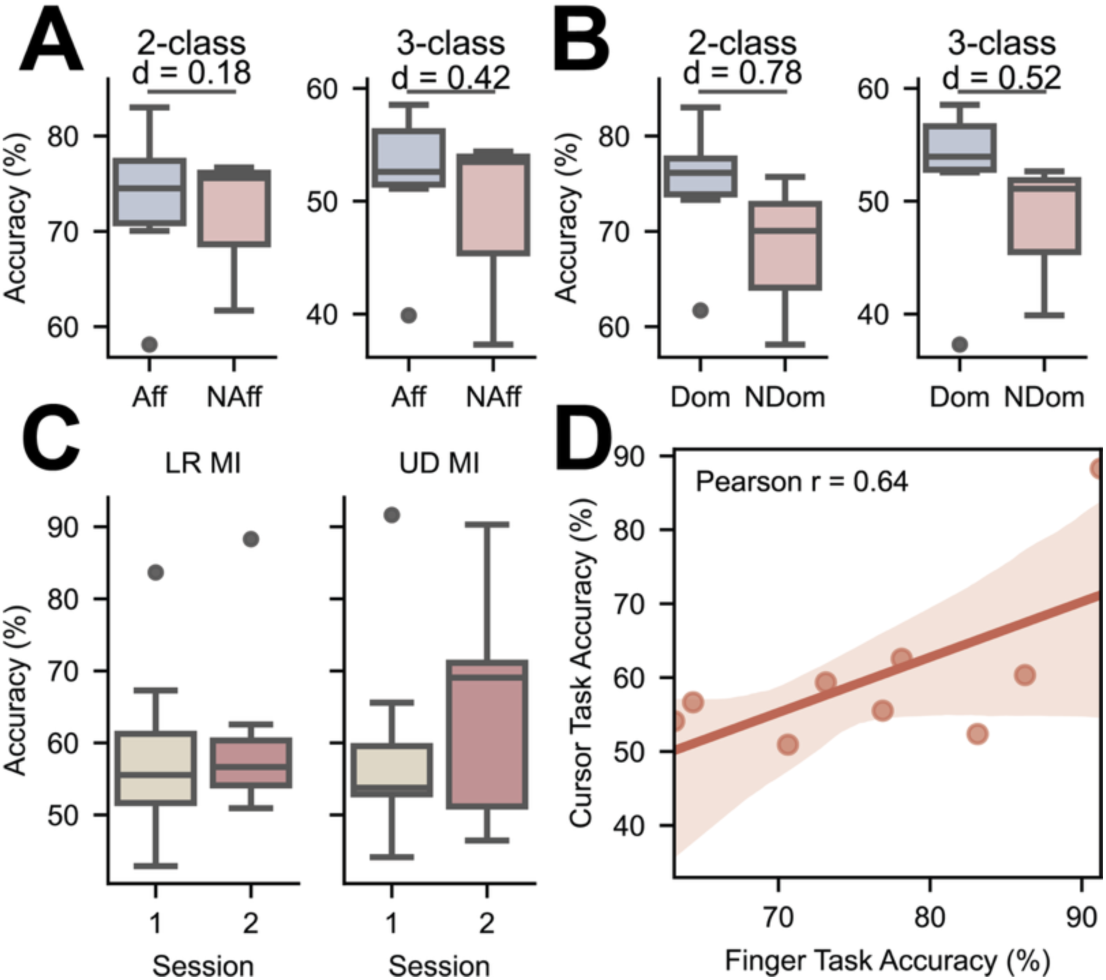
Subgroup analysis of online finger MI control performance and its relationship to limb-level MI decoding. (**A**) Comparison of 2-finger and 3-finger online decoding performance between stroke survivors who performed MI control using their affected hand (*n* = 6) and those using their non-affected hand (*n* = 3). (**B**) Comparison of 2-finger and 3-finger online decoding performance between stroke survivors who performed MI control using their dominant hand (*n* = 6) and those using their non-dominant hand (*n* = 3). (**C**) Online performance in the limb-level cursor MI training task across two sessions. (**D**) Correlation between online finger-level MI decoding accuracy and cursor-task performance (*n* = 9). The solid line represents the linear regression fit, and the shaded region denotes the 95% confidence interval.

**S3 Fig.**
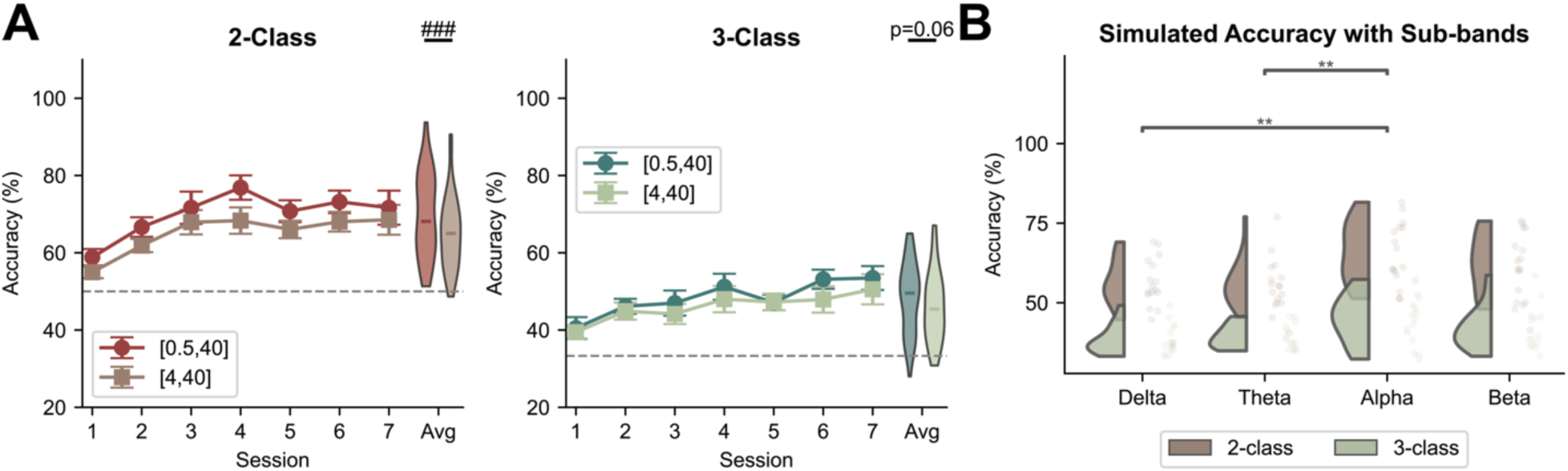
Offline analysis of low-frequency contributions. (**A**) Simulated offline base decoding accuracy using EEG inputs filtered at 0.5–40 Hz and 4–40 Hz for 2-finger and 3-finger MI across seven online sessions (*n* = 9). The grey dashed line represents the chance level, and error bars indicate the standard error. The performance distribution across all sessions is shown on the right. The center lines indicate the median value. The boxes extend from the lower quartile to the upper quartile, and the lines indicate 1.5 times the interquartile range. Statistical analysis was performed using a two-way repeated-measures ANOVA with main effects of session and bandpass filtering setting. Significant main effects of bandpass are indicated (### if *p* < 0.001). (**B**) Simulated decoding accuracy using EEG signals filtered into individual frequency bands for the healthy control group (delta, theta, alpha, and beta) (*n* = 16). Statistical significance was observed for the main effect of frequency bands in a two-way ANOVA. Significance stars indicate post hoc pairwise comparison results using an FDR-corrected two-tailed Wilcoxon signed-rank test (** if *p* < 0.01).

**S4 Fig.**
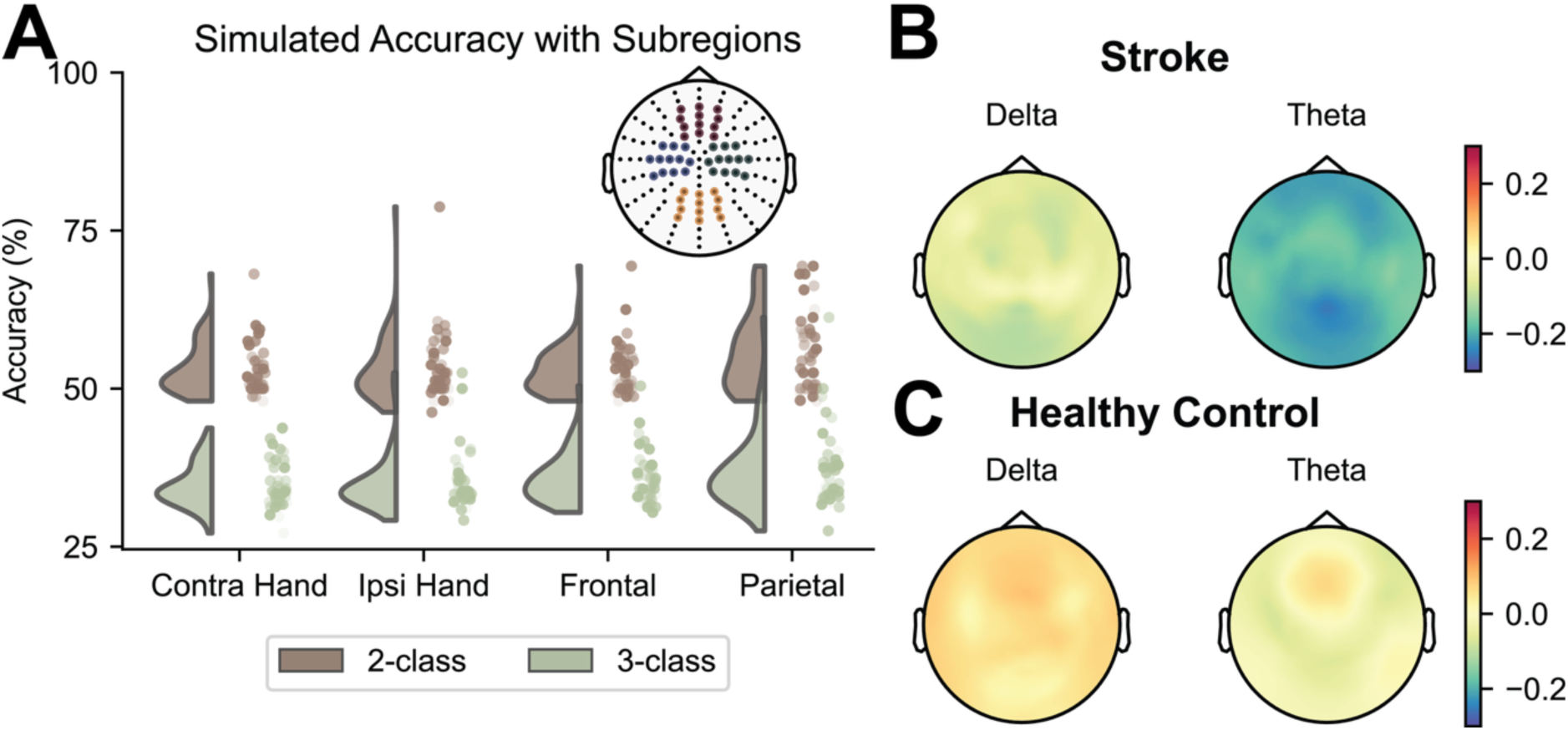
Offline analysis of cortical subregion contributions and task-induced electrophysiologies. **(A)** Simulated decoding accuracy using EEG signals within individual cortical subregions (*n* = 9). The channels within the contralateral hand (Contra Hand), ipsilateral hand (Ipsi Hand), frontal, and parietal regions are colored in blue, green, red, and yellow, respectively. **(B** and **C)** Group-averaged topographical maps of delta-band (0.5-4 Hz) and theta-band (4-8 Hz) ERD during online MI control for the stroke group (**B,** *n* = 9) and the healthy control group (**C,** *n* = 16).

**S5 Fig.**
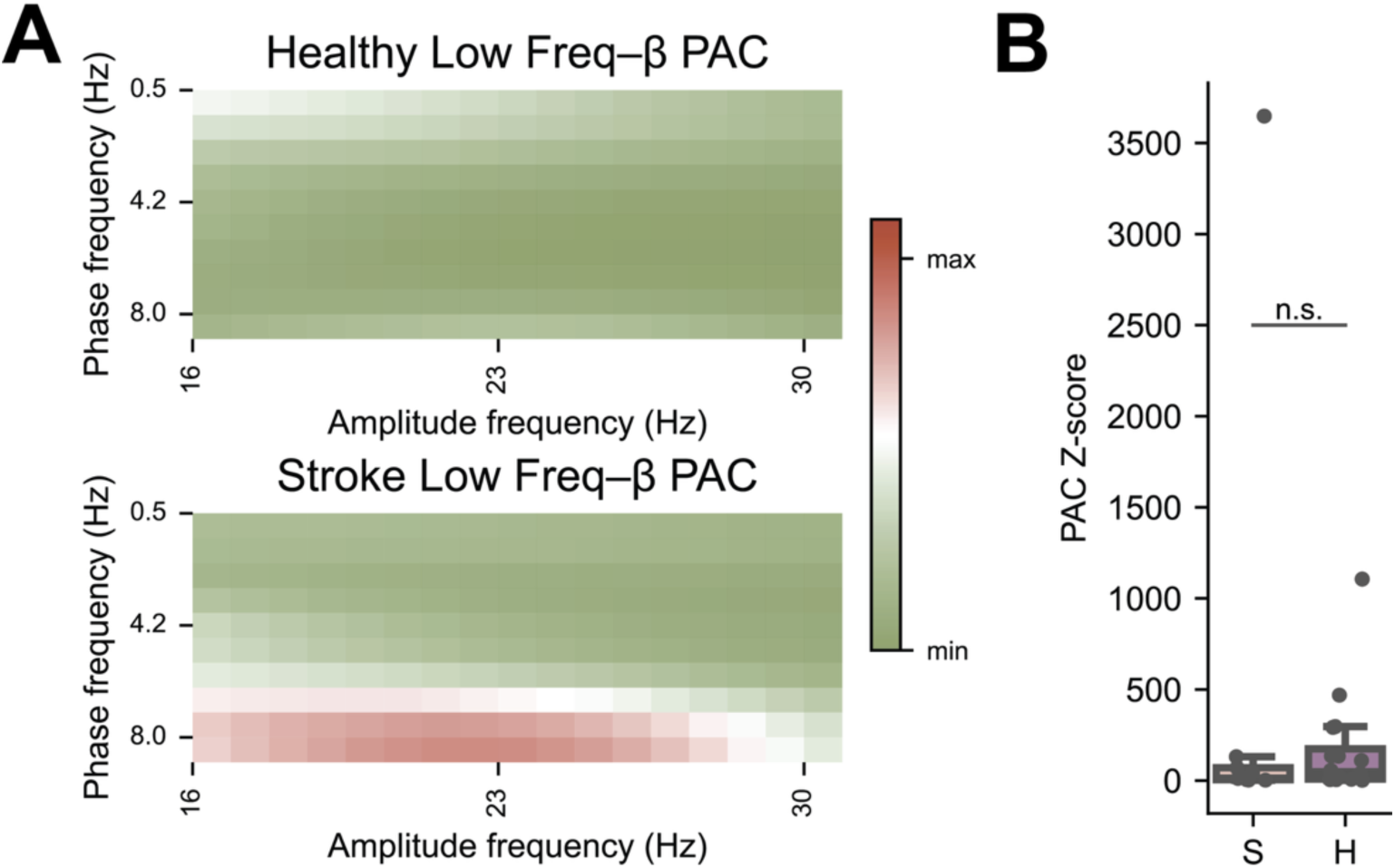
Cross-frequency correlation in stroke survivors. (**A**) The phase-amplitude coupling (PAC) of the stroke and healthy group between low frequency activities (0.5 – 8 Hz) and beta activities over the contralateral sensorimotor cortex (the ipsilesional sensorimotor cortex for stroke survivors). PAC values were normalized to the maximum value observed across the two groups for visualization. (**B**) Comparison of PAC z-scores between groups. Z-scores were computed based on surrogate distributions generated using 200 permutations. Between-group differences were assessed using two-tailed Mann–Whitney U tests (n.s. if no statistical significance is found).

**S6 Fig.**
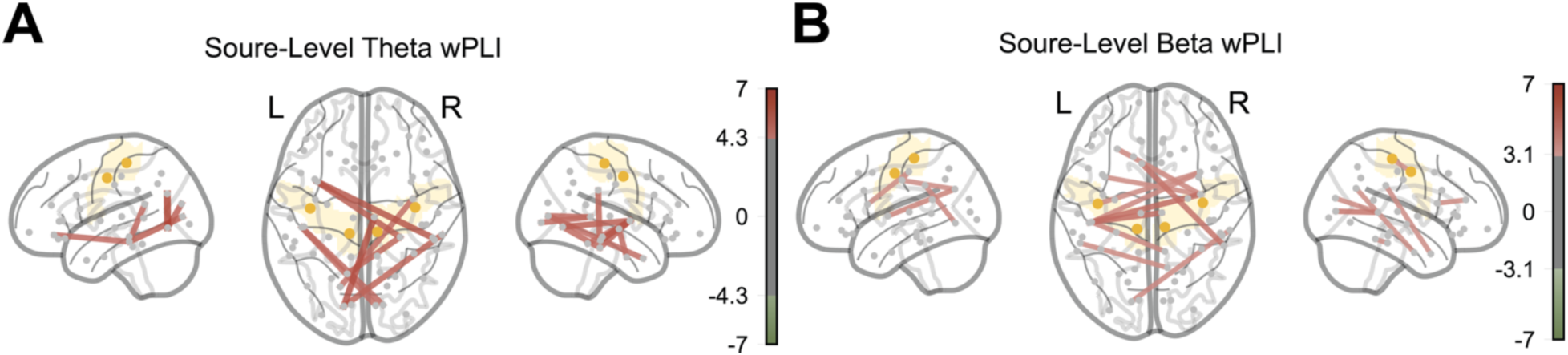
Group-level differences in theta-band (A) and beta-band (B) weighted Phase Lag Index (wPLI) at the source level, expressed as z-scores relative to healthy controls. Functional connections between the atlas with absolute z-scores exceeding the 99th percentile threshold are displayed, with edge color and thickness indicating the sign and magnitude of the z-score.

## Notes

### Competing Interest Statement

YD and BH are co-inventors of a pending patent application related to robotic finger BCI technique used in this work.

